# The rapid decline in interaural-time-difference sensitivity for pure tones can be explained by peripheral filtering

**DOI:** 10.1101/2023.08.04.551950

**Authors:** Matthew J. Goupell, G. Christopher Stecker, Brittany T. Williams, Anhelina Bilokon, Daniel J. Tollin

**Author notes:** Corresponding Author: Matthew J. Goupell Department of Hearing and Speech Sciences University of Maryland College Park, MD 20742.

## Abstract

**Purpose:** The interaural time difference (ITD) is a primary horizontal-plane sound localization cue computed in the auditory brainstem. ITDs are accessible in the temporal fine structure of pure tones with a frequency of no higher than about 1400 Hz. Explaining how listeners’ ITD sensitivity transitions from very best sensitivity near 700 Hz to impossible to detect within 1 octave currently lacks a fully compelling physiological explanation. Here, it was hypothesized that the rapid decline in ITD sensitivity is dictated not by a central neural limitation but by initial peripheral sound encoding, specifically, the low-frequency (apical) edge of the cochlear excitation pattern produced by a pure tone.

**Methods:** ITD sensitivity was measured in 16 normal-hearing listeners as a joint function of frequency (900-1500 Hz) and level (10-50 dB sensation level).

**Results:** Performance decreased with increasing frequency and decreasing sound level. The slope of performance decline was 90 dB/octave, consistent with the low-frequency slope of the cochlear excitation pattern.

**Conclusion:** Fine-structure ITD sensitivity near 1400 Hz may be conveyed primarily by “off-frequency” activation of neurons tuned to lower frequencies near 700 Hz. Physiologically, this could be realized by having neurons sensitive to fine-structure ITD up to only about 700 Hz. A more extreme model would have only a single narrow channel near 700 Hz that conveys fine-structure ITDs. Such a model is a major simplification and departure from the classic formulation of the binaural display, which consists of a matrix of neurons tuned to a wide range of relevant frequencies and ITDs.

## INTRODUCTION

Humans can precisely localize sounds in the horizontal plane. When a sound is located away from the midline, the extra distance that the sound travels to the far ear generates an interaural time difference (ITD) (Hartmann, 2021). There is an obvious central physiological substrate for ITD processing in the medial superior olivary complex of the mammalian auditory brainstem, which consists of neurons that primarily respond to low frequencies and receive bilateral frequency-matched inputs from the anteroventral cochlear nucleus from cells that have short-latency and secure synaptic connections with superior temporal precision (Müller, 1990; Blanks et al., 2007; Yin et al., 2019; Litovsky et al., 2021).

The most important ITD cues for sound localization occur at frequencies below 1500 Hz (Wightman and Kistler, 1992; Macpherson and Middlebrooks, 2002) and are conveyed in the acoustic waveform’s temporal fine structure, the temporal aspects of the rapid variation in pressure that the auditory nerve can precisely encode. There are two notable features to ITD frequency dependence below 1500 Hz. First, the best sensitivity in ITD discrimination for pure tones is exquisitely good, on the order of 10 µs. This occurs for pure tones near 700 Hz (Klumpp and Eady, 1956; Zwislocki and Feldman, 1956; Thavam and Dietz, 2019; Klug and Dietz, 2022), which coincides with the often-called ITD frequency “dominant region” (i.e., a region of very high relative weight) for lateralization of narrowband noises, transients, and complex tones that exceed a single peripheral cochlear filter width (Stern et al., 1988; Shackleton et al., 1992; Tollin and Henning, 1999; Folkerts and Stecker, 2022; Goupell and Bilokon, 2022). Second, ITD discrimination sensitivity for pure tones declines at a remarkable slope previously estimated to be at least 46-78 dB/octave as frequency is increased from 1200 to 1400 Hz (Brughera et al., 2013; Klug and Dietz, 2022). This is much steeper than the decline in temporal information or phase locking (i.e., the ability to generate action potentials during a preferred phase of the waveform) conveyed by auditory nerve (12 dB/octave; Klug et al., 2023), the spherical bushy cells of the anteroventral cochlear nucleus (<12 dB/octave; Joris et al., 1994; Klug and Dietz, 2022), or model MSO neurons (20 dB/octave; Brughera et al., 2013; Klug and Dietz, 2022). This makes the decline in neural phase locking between the periphery and brainstem an insufficient explanation for the behavioral phenomenon. The question posed in this study is whether the upper frequency limit of fine-structure ITD processing might be related to the frequency of best ITD processing and the dominant region. This could be a result of some unique aspect of binaural processing at 700 Hz that allows for the superior conveyance of fine-structure ITD.

Most studies that investigate the frequency dependence of ITD discrimination sensitivity present stimuli at a fixed level, for example 65 dB SPL (Klumpp and Eady, 1956) or 70 dB SPL (Brughera et al., 2013; Klug and Dietz, 2022), comparable to the level of conversational speech. Lower level sounds (e.g., 10 dB sensation level, SL) are perceived relatively nearer the midline of the head (Ihlefeld et al., 2019) and have worse ITD discrimination thresholds (Dietz et al., 2013). For low-frequency pure tones, Zwislocki and Feldman (1956) measured ITD discrimination thresholds in up to six normal-hearing listeners as a function of level (10-110 dB SL) and frequency (250, 500, 1000, and 1250 Hz). ITD performance decreased as the frequency increased from 1000 to 1250 Hz, and as the level decreased from 70 to 10 dB SL. At 1250 Hz and 10 dB SL, the ITD discrimination threshold was not measurable, which places the upper frequency limit of fine-structure ITD processing at a frequency that is much lower than the typically reported 1400 Hz for higher level tones (Klug and Dietz, 2022).

Few other stages of the auditory system that shows an incredibly steep >>20 dB/octave slope of decline. One other obvious location that demonstrates such a steep rate is at the auditory periphery in the cochlea, where there is a high-resolution frequency decomposition of sound. In the cochlea, sound causes frequency-specific pattern of displacement along the length of the basilar membrane, such that successive segments behave effectively like a bank of narrowly tuned bandpass filters (Oxenham and Shera, 2003). The transformation of basilar membrane displacement to an “excitation pattern” is a way to represent the transformation of a physical stimulus to an internal spectral stimulus representation (Moore and Glasberg, 1983). The excitation pattern is asymmetrical across frequency, specifically featuring sharp low-frequency (apical) and shallow high-frequency (basal) tails. Critically, the low-frequency (apical) edge of the excitation pattern generated by a tone passing through one of these peripheral bandpass auditory filters is approximately 100 dB/octave at 1000 Hz (Oxenham and Shera, 2003; Whiteford et al., 2020). This means that the excitation pattern of a tone extends and induces activity in neurons tuned to lower frequencies (apically).

Consider the possibility that the steep slope of the low-frequency (apical) edge of the excitation pattern and the rapid decline in ITD sensitivity could be related. First, assume that the auditory filter near 700 Hz is the best and highest frequency filter that can convey temporal fine structure ITD. If this is the case, then increasing the stimulus frequency higher than 700 Hz decreases the energy in this best auditory filter. This would directly impair ITD discrimination sensitivity, which decreases with decreasing level (Zwislocki and Feldman, 1956; Dietz et al., 2013). This is important because this activity would decrease rapidly with increasing tone frequency and decreasing tone level (Fig. 1). At some frequency near 1400 Hz, the activity of neurons near 700 Hz would cease altogether regardless of the tone level (even at a high level of, for example, 70 dB SPL) due to the exceptionally steep excitation pattern. That expectation is consistent with the abrupt cessation of the ability to use fine-structure ITDs near 1400 Hz.

**Fig. 1:**
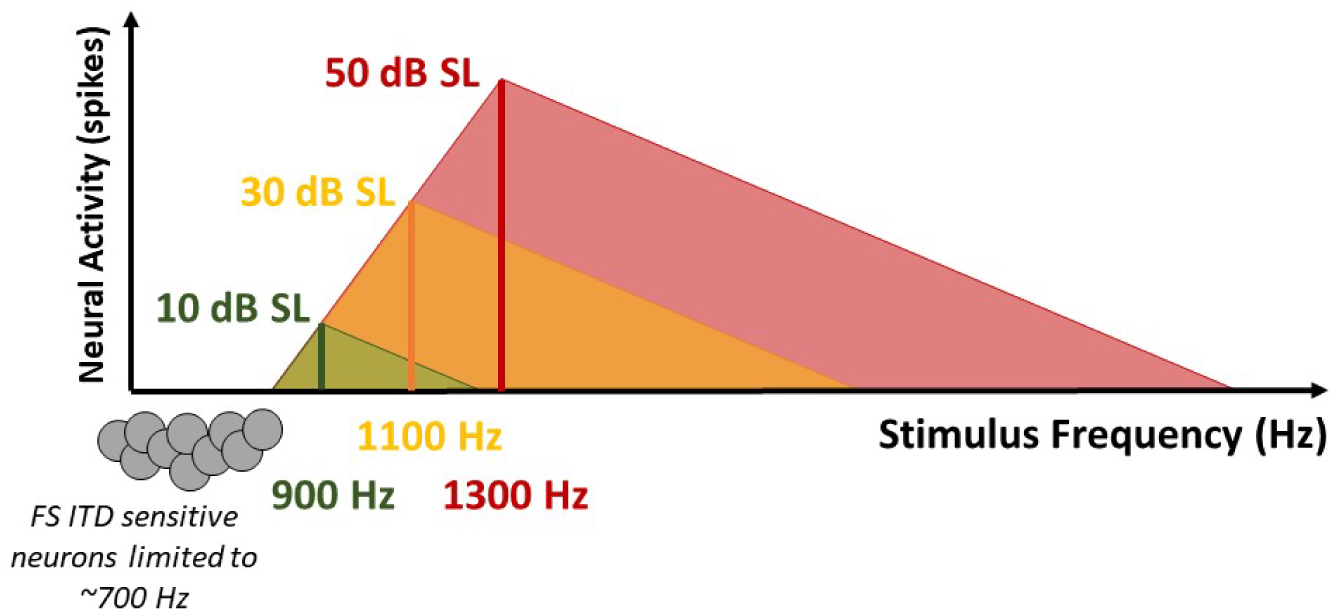
Illustration of the transformation of input acoustic tones into their effective neural spectral representations [i.e., their excitation patterns, which have a steep low-frequency (apical) slope and shallow high-frequency (basal) slope] and their relationship to the experimental hypothesis. If fine-structure ITD is conveyed by neurons that are limited to a highest frequency of about 700 Hz, decreasing the stimulus level will decrease the highest frequency that demonstrates fine-structure ITD sensitivity. In this example, the excitation pattern of a 1300-Hz 50-dB-SL tone (red) excites the lower-frequency fine-structure ITD-sensitive neurons. Decreasing the stimulus level lowers the maximum frequency of a tone that excites these neurons because it limits the extent of the excitation pattern. The tone needs to be 1100 Hz for 30 dB SL (yellow) and 900 Hz for 10 dB SL (green) for their excitation patterns to excite the lower-frequency (700 Hz) fine-structure ITD-sensitive neurons. The rate of performance decline should change by approximately the same slope as the low-frequency (apical) edge of the cochlear excitation pattern, about 100 dB/octave (Oxenham and Shera, 2003).

Such a relationship can be better understood if the upper frequency limit of fine- structure ITD sensitivity was investigated as a function of level. Such measurements, however, have not been made with adequate resolution to quantify the slope of ITD sensitivity decline. Therefore, this study measured this upper frequency limit as a function of sound level. It was hypothesized that the upper frequency limit of fine-structure ITD processing decreases with decreasing level with a slope near 100 dB/octave, suggesting it is a consequence of cochlear filtering, the low-frequency edge of the cochlear excitation pattern. Through the confirmation of this hypothesis, we have shown that these data can be explained by the cochlear filtering, and that it is possible that fine-structure ITD is conveyed through responses of neurons limited to only 700 Hz and possibly from a single dominant channel, contradicting the classic formulation of the binaural display.

## METHODS

### Listeners and Equipment

Sixteen young to middle-aged (mean=30.1 yrs, range=20-51 yrs) normal-hearing listeners (10 female) were tested in this study. Hearing thresholds were measured at octave frequencies from 0.25-8 kHz. Hearing thresholds were ≤20 dB hearing level (HL) for all listeners except C1, who had slightly higher thresholds ≥6 kHz. Interaural asymmetry in hearing thresholds was ≤10 dB except it was 15 dB at 1000 Hz for listener M5. Data from the authors (listeners M1, B1, B2, and C1) were included.

The testing was performed at three locations (Maryland, Nebraska, and Colorado), three to seven listeners at each location. The same testing software and type of circumaural headphones (HD280 Pro, Sennheiser; Hannover, Germany) was used at each location. Stimuli were generated and the experiment was conducted on a personal computer using MATLAB (the Mathworks; Natick, MA). Procedures were reviewed and approved by the University of Maryland, Boys Town National Research Hospital in Nebraska, and the University of Colorado School of Medicine Multiple Institutional Review Boards for use of human subjects.

In Maryland, testing was performed in a double-walled sound booth (IAC; North Aurora, IL). Stimuli were presented to listeners via an external sound card (Sound Blaster PLAY! 3, Creative Labs; Milpitas, California). In Nebraska, testing was performed in a double-walled electromagnetically shielded sound booth (ETS-Lindgren; Cedar Park, TX). Stimuli were presented to listeners via USB audio interface (RME Fireface UFX; Haimhausen, Germany). In Colorado, testing was performed in a single-walled custom-built sound booth. Stimuli were presented to listeners via an external digital-to-analog converter (Schitt Modi, Schitt Audio; Newhall, CA) and headphone amplifier (Schitt Magni 3+, Schitt Audio; Newhall, CA).

### Stimuli

The stimuli were pure tones that had a frequency of 900, 1000, 1100, 1200, 1300, 1400, or 1500 Hz. For audiometric threshold measurements, tones were presented diotically. For ITD discrimination testing, tones started with the left channel in sine (0°) phase and the right channel had the same phase or a phase that yielded ±167-µs ITDs. The ITD was applied in only the temporal fine structure and there was a slow diotic envelope was intended to minimize the contribution of an envelope ITD (Stecker and Bibee, 2014). After the diotic raised-cosine temporal ramp was applied, the stimuli had a 100-ms rise-fall time and 50-ms full on duration. In other words, the total duration of the tone was 250 ms. A visualization of the stimuli can be seen in Fig. 2. The tone amplitude was 10-, 20-, 30-, or 50-dB SL in different conditions.

**Fig. 2:**
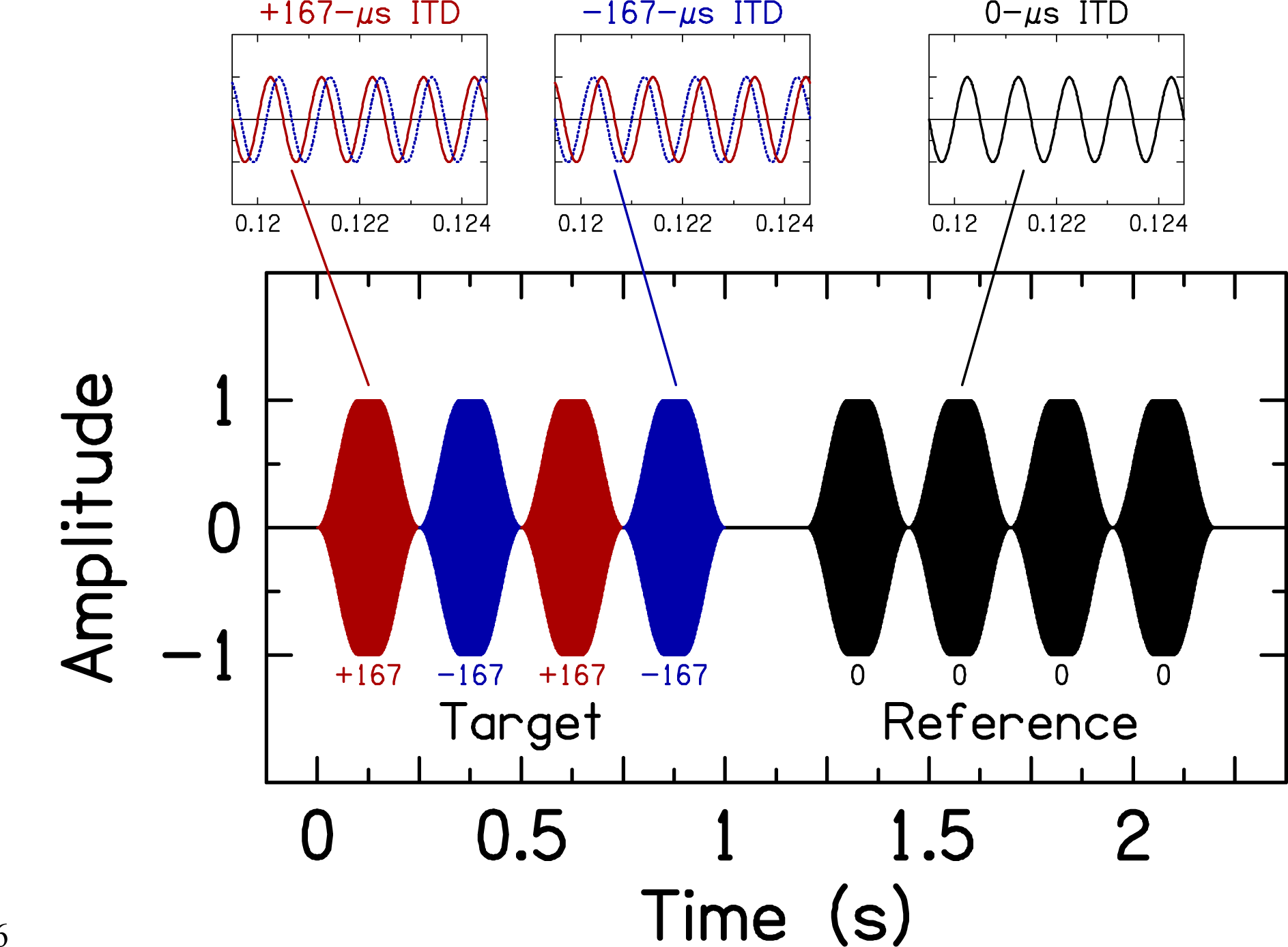
Stimulus waveform for a 1000-Hz tone. The first interval is the target interval, which contains four tone bursts where the ITD alternates between right (red) and left (blue) leading 167-µs ITDs. The second interval is the reference interval, which contains four tone bursts where the ITD is zero. Small panels above the large panel show the waveforms with a smaller time range to see the ITD more clearly. For stimulus portions with a non-zero ITD, solid red traces represent the right channel, dotted blue traces represent the left channel. For stimulus portions with a zero ITD, both traces overlap and are solid black.

### Procedure

#### Pure-Tone Audiometric Threshold Measurements

Pure tones were presented diotically. An experimenter controlled the timing of presentation and level of the stimulus by using buttons that could adjust the level by 0, ±1, ±2, or ±6 dB. Listeners indicated when they heard a tone. Experimenters adjusted the sound level above and below the level that produced threshold (transition from response to no response) at least twice to determine a final threshold level (TL), which defines 0 dB SL, at each test frequency. The frequencies were tested in increasing order. The threshold at each frequency was tested twice for most listeners. If the two thresholds differed by >5 dB, additional threshold levels were measured and the average of the last two measurements were used. Across-frequency averaging was implemented to smooth out small variations in TL during later periods of testing (listeners M6, B4, B5, and B6): the average TL at each frequency was computed as the average of levels from the target and adjacent frequencies, rounded to the nearest integer. For example, the TL at 1200 Hz was based on the six measurements from 1100, 1200, and 1300 Hz. Adjustments due to averaging were small (generally, less than 2 dB; on average <1 dB over all listeners and frequencies) and yielded comparable results to unsmoothed TLs used for other listeners.

#### Left-right ITD Discrimination

Listeners performed a two-interval forced-choice discrimination task, where each interval contained four tone bursts, which was based on the approach used by Füllgrabe and Moore (2018). Figure 2 shows an example stimulus. Each interval presented a sequence of four 250-ms tone bursts. In the reference interval, all four tone bursts had zero ITD. In the target interval, the ITD alternated twice between right and left leading ±167- µs ITDs (i.e., right-left-right-left). The ITD changed sign at the envelope minimum to avoid a waveform discontinuity (e.g., Undurraga et al., 2016). At each frequency, a constant ITD was used instead of a constant IPD because lateralization extent–the likely cue used to perform the left-right discrimination task–appears to depend on ITD not IPD (e.g., Zhang and Hartmann, 2006). Moreover, medial superior olive neurons encode ITDs (e.g., Pena and Konishi, 2000). The ITD magnitude was chosen because it corresponds to one-quarter of the stimulus period at 1500 Hz, which was the highest frequency tested in this study. This ITD guarantees a monotonic psychometric function up to 1500 Hz because it avoids the descending portion of the cyclical psychometric function (Klug and Dietz, 2022).

There was a 200-ms silent interval between the target and reference intervals. The order of the intervals was randomized with a probability of 50%. The task of the listener was to identify which interval contained the target alternating ITD.

The task was self-paced because listeners initiated each trial. After the stimulus was presented, the listener indicated the target interval by pressing a button on the screen. Correct answer feedback was provided after each trial, which helped keep the listener attentive during testing.

Pre-experiment training was performed to ensure the listener understood the task and had no apparent problems with the task. Listeners were initially presented with 10 trials at 500 Hz and 70 dB SL. Training in 10-trial blocks continued until performance was 100% correct. Almost all the listeners needed only the initial 10 practice trials to proceed.

For the main experiment, a method of constant stimuli was performed where all combinations of frequency and level were presented in random order, consisting of seven frequencies (900–1500 Hz in 100-Hz steps) and four levels (10, 20, 30, and 50 dB SL). Therefore, there were 112 trials per block (7 frequencies×4 level×4 trials), and each block took ∼10 minutes to complete. Testing breaks were provided between blocks as needed. Listeners performed 60 trials per condition (i.e., the specific combination of frequency and level). Three listeners were provided less than 60 trials (minimum of 56 trials) for at least one condition and four listeners were provided more than 60 trials (maximum of 83 trials) for at least one condition because of computer or experimenter error. The experiment took ∼3 hours to complete.

## RESULTS

### The upper frequency limit of fine-structure ITD sensitivity decreases from 1400 to 900 Hz with decreasing sound level

Individual ITD discrimination data are shown in Fig. 3 for all 16 normal-hearing listeners. The average performance calculated across all tested conditions is reported in each panel. While there is variability in the overall performance, the patterns in the data are consistent across listeners and testing sites. Two listeners (M6 and B3) had great difficulty with this task, with an average performance <60%. These two listeners were removed from subsequent group analyses. For all other listeners, ITD discrimination performance decreased systematically with increasing frequency and decreasing sound level.

**Fig. 3:**
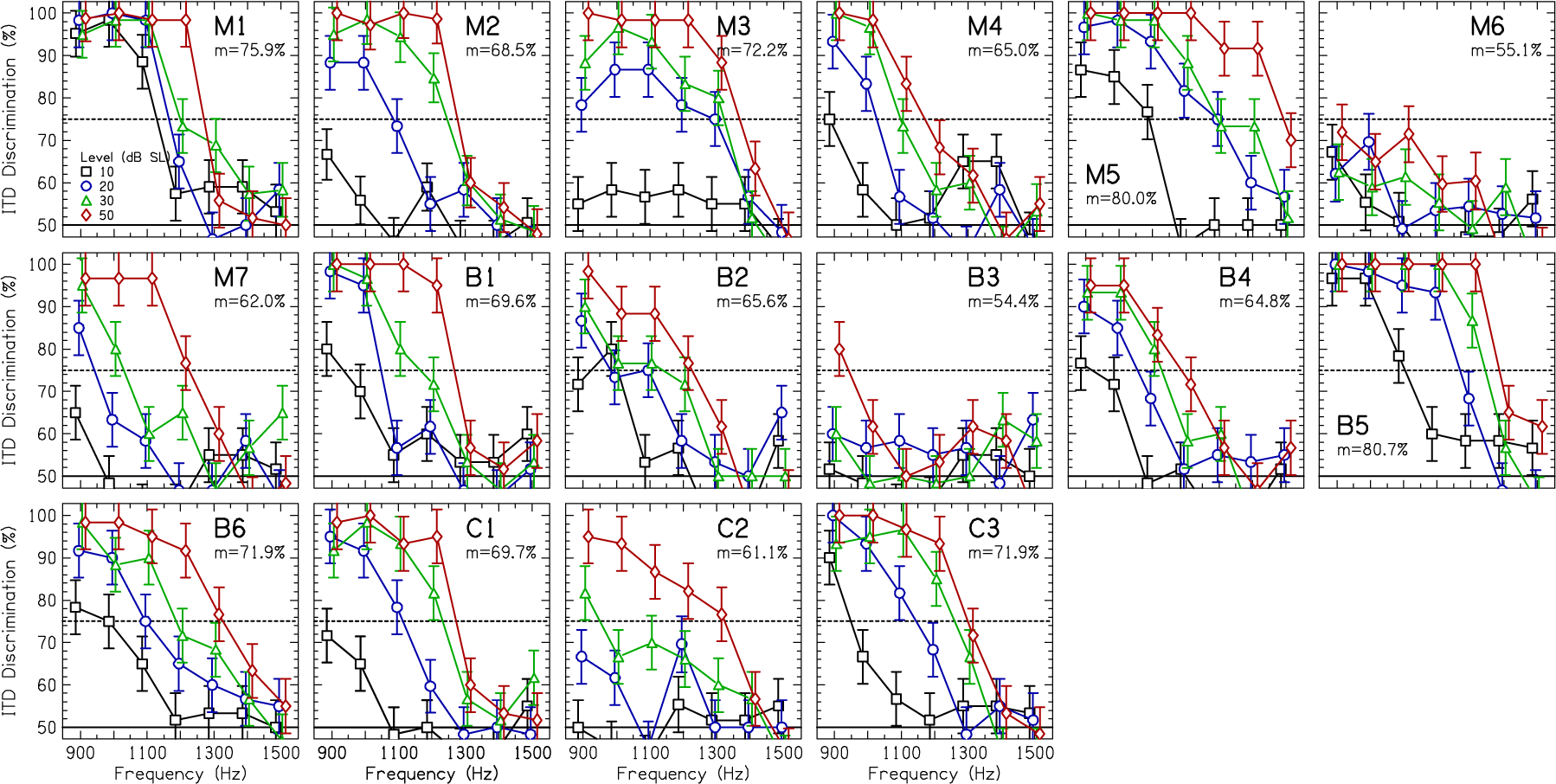
Percent correct as a function of frequency and level (in dB SL) for 16 listeners. Error bars show 95% confidence intervals. The dashed horizontal line denotes 75% correct. The average percent correct over all conditions is reported in each panel.

The average ITD discrimination data are shown in Fig. 4(A) for 14 listeners. A three-way repeated-measures analysis of variance was performed on ITD discrimination performance transformed to rationalized arcsine units (Studebaker, 1985) with factors frequency (7 levels: 900-1500 Hz), stimulus level (4 levels: 10, 20, 30, 50 dB SL), and testing location (3 levels: Maryland, Nebraska, Colorado). The Greenhouse-Geisser correction was used to address sphericity assumption violations. There were significant effects of frequency [F(2.8,5.6)=73.7, *p*<0.0001, *η*_*p*_^2^=0.87] and stimulus level [F(1.6,3.2)=99.4, *p*<0.0001, *η*_*p*_^2^=0.90]. There was a significant frequency×stimulus level interaction [F(4.4,8.8)=13.9, *p*<0.0001, *η*_*p*_^2^=0.56]. The effect of testing location was not significant [F(2,11)=0.265, *p*=0.772, *η*_*p*_^2^=0.05], and there were no significant interactions with testing location (*p*>0.05 for all).

**Fig. 4:**
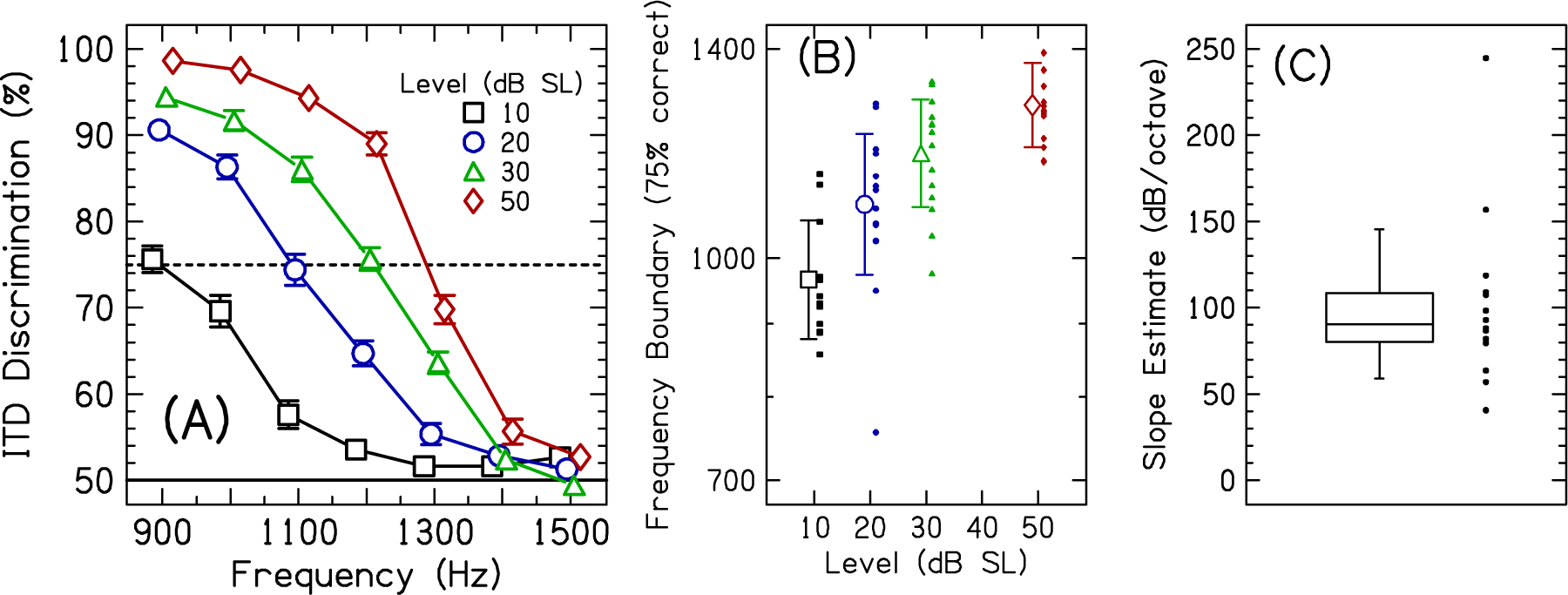
(A) Percent correct as a function of frequency and sensation level averaged over 14 listeners. Error bars represent ±1 standard error. Some conditions had error bars that were smaller than the symbol. The dashed horizontal line denotes 75% correct. (B) Frequency boundaries that yield 75% correct ITD discrimination performance. Small closed symbols show individual data. Large open symbols show the data averaged over listeners. The error bars represent ±1 standard deviation. (C) Linear slope estimates between sound level (in dB SL) and frequency (in octaves). Small closed symbols show the slope for the individual listeners. The box length represents the interquartile range, and the median is shown by the horizontal line within the box. Whiskers extend to 10^th^ and 90^th^ percentiles.

The upper frequency limit of fine-structure ITD processing (defined as 75% correct ITD discrimination performance) was found for each presentation level and for each listener by fitting the data with a logistic regression (Wichmann and Hill, 2001). The frequency boundaries are shown in Fig. 4(B). These data generally agree with others that show that the upper frequency limit of fine-structure ITD processing is near 1400 Hz for sound levels above 50 dB SPL (Klumpp and Eady, 1956; Zwislocki and Feldman, 1956; Brughera et al., 2013; Klug and Dietz, 2022). Across listeners, the frequency that yielded 75% correct ITD discrimination was 1264.8±95.4 Hz (range=1168–1483) for 50 dB SL tones. For better comparison to past studies estimating the upper frequency boundary at higher levels, the data in Fig. 4(B) were fit by a three-parameter exponential rise to maximum function. The best fitting equation (r=0.74, *p*<0.0001) was *y* = 775.9 + 587.6 × (1 − *e*^−0.039*x*^), where *y* is the frequency boundary and *x* is the sensation level.

Evaluating this function at 70 dB SL yielded an estimated upper frequency boundary of 1326 Hz with a 95% confidence interval from 1212 to 1437 Hz. This confidence interval encompasses the upper frequency boundary measured in other studies (Klumpp and Eady, 1956; Zwislocki and Feldman, 1956; Brughera et al., 2013; Klug and Dietz, 2022).

Any other small discrepancies in the upper frequency limit of fine-structure ITD processing between the current study and others can be explained by methodological differences. For example, our stimuli had slow diotic ramps minimizing contributions from onset ITD (Stecker and Bibee, 2014). We also tested more listeners than previous studies, and thus may have included more listeners that were not operating at a point near the absolute limits of human binaural processing, including two that were middle aged (Ross et al., 2007; Füllgrabe and Moore, 2018). Similarly, Klug and Dietz (2022) and Best and Swaminathan (2019) tested a relatively large number of normal-hearing listeners (n=9), and reported relatively larger ITD thresholds compared to some studies with relatively smaller (potentially more selective) samples. It may also be that more extensive training would improve ITD discrimination performance to the levels of the best listeners in this study (Wright and Fitzgerald, 2001), including the two who had <60% total correct responses and were omitted from the average data and slope calculations (described below).

These data also agree with other studies that show declines in ITD discrimination performance with decreasing sound level (Zwislocki and Feldman, 1956; Dietz et al., 2013; Best and Swaminathan, 2019). This decrease in performance with decreasing level causes the 75% correct performance point to shift from about 1400 to 900 Hz [Fig. 4(B)]. It is notable that changing the stimulus level from 50 to 30 dB SL produces a marked change in the upper frequency limit given that both stimuli are clearly audible. This result is also difficult to reconcile with a phase-locking explanation, which reaches a maximum within ∼20 dB of neural threshold (Palmer and Russell, 1986). In other words, both the rate of fine-structure ITD discrimination performance decline across frequency for a fixed level and the change across level for a fixed frequency is difficult to reconcile using current binaural and physiological models. Ultimately, it appears that it is important to account for the stimulus sound pressure level when considering the upper frequency limit of fine- structure ITD processing and future binaural modeling efforts that account for level effects could advance the field.

### The rate of fine-structure ITD sensitivity decline is 90 dB/octave, which can be explained by peripheral cochlear filtering

A linear fit between the sound level (in dB SL) and the frequency boundary (in octaves relative to 1000 Hz) determined the slope of the ITD performance change. The 10 dB SL point was omitted from the calculation for 2 of the 14 listeners (M3 and C2) because they did not have an estimate of the frequency boundary at 75% correct (i.e., they were guessing in this condition). The resulting fit slopes are shown in Fig. 4(C). The median slope was 90.5 dB/octave (range=40.7–244.6). The linear fits explained a median of 88% in the variance of the data points (range=74–99.8).

The estimated slope of 90.5 dB/octave could be considered the conservative lower boundary. Had the stimuli used constant IPD as a function of frequency, there would have been increasingly large ITDs with decreasing frequency from 1500 Hz. This means that performance would have increased at a more rapid rate and the estimated slope could have been even steeper.

The most important addition to our knowledge that results from this study is that the slope of ITD discrimination performance decline was measured to be 90 dB/octave [Fig. 4(C)]. Such a steep slope limits the possible explanations for the physiological mechanisms that are involved in the upper frequency limit of fine-structure ITD processing. For example, Klug *et al*. showed that phase locking in the auditory nerve across animal models declines at approximately 12 dB/octave (Klug et al., 2023). Phase locking at the brainstem, anteroventral cochlear nucleus and medial superior olive, also do not seem to decline sufficiently rapidly (Klug and Dietz, 2022). Therefore, the decline in neural phase locking at and before the brainstem by itself cannot account for these data because the slopes are so different. In contrast, one well-known stage of the auditory system that does have such a steep slope is the low-frequency edge of the cochlear excitation pattern, which is approximately 100 dB/octave near 1000 Hz (Oxenham and Shera, 2003; Whiteford et al., 2020).

## DISCUSSION

The upper frequency limit of human fine-structure ITD processing was determined by perceptual ITD discrimination measurements that sampled frequency and sound level with a high resolution. It was found that ITD discrimination sensitivity decreases with decreasing level with a slope of 90 dB/octave.

Our interpretation of the data is that an auditory filter near 700 Hz is the highest frequency filter that can convey temporal fine structure ITD. Best ITD discrimination sensitivity occurs for a 700-Hz tone and increasing the stimulus frequency higher than 700 Hz decreases the energy in this auditory filter. ITD discrimination sensitivity decreases with decreasing level (Zwislocki and Feldman, 1956; Dietz et al., 2013), such that percent correct is directly related to effective stimulus intensity. Therefore, the slope analysis yields the rate at which the energy in this auditory filter decreases. Given that the measured slope is incredibly steep for the auditory system, 90 dB/octave, we conclude that it is possible that this slope is a result of peripheral filtering [i.e., the low-frequency (apical) edge of the cochlear excitation pattern; Fig. 1], rather than the decay in phase-locking at the level of the auditory nerve or brainstem (Klug and Dietz, 2022; Klug et al., 2023).

If the peripheral cochlear filtering interpretation of these data is correct, it has a further important corollary. It suggests that ITD sensitivity for stimulus frequencies between 1000-1500 Hz are primarily driven by lower-frequency neurons, such as those tuned between 500-1000 Hz, in the region where the best ITD sensitivity is observed (Klumpp and Eady, 1956) and location of the ITD frequency dominant region (Fig. 1) (Folkerts and Stecker, 2022). It could be that fine-structure ITD sensitive neurons extend to all frequencies below 700 Hz. Alternatively, there could be a narrow band of neurons near 700 Hz in humans that is responsible for conveying fine-structure ITD, which in turn yields the exquisite sensitivity to fine-structure ITDs and carries most of the localization weight for broadband stimuli. Indeed, examination of the functional form of ITD sensitivity as a function of frequency [see Fig. 9 in Klug and Dietz (2022)] resembles the neural response area of a high-level tone (i.e., a shallow low-frequency portion, a minimum at 700 Hz, a sharp high-frequency portion; the inverse of the excitation patterns shown in Fig. 1), which would suggest the possibility of a single dominant channel for fine-structure ITD processing. To the extent that neurons tuned to 1000-1500 Hz contribute to fine-structure ITD sensitivity, they are clearly less important than those between 500-1000 Hz because the observed transition slope does not extend near the high-frequency edge of their tuning, which would be 2 kHz or higher. They may, in fact, not contribute at all. It is possilbe that the dominant region neurons are highly weighted even for single pure tones.

The individual variability in ITD discrimination sensitivity (Fig. 3) appears to be consistent with range of binaural performance in other reports with ∼10 listeners. For example, Best and Swaminathan (2019, Fig. 1) reported ITD thresholds at 750 Hz to vary by a factor of 10 or more (20-200 µs in nine normal-hearing listeners, 20-300 µs or more in nine hearing-impaired listeners). Klug and Dietz (2022, Fig. 7) reported ITD thresholds at 800 Hz that varied by a factor of five (20-100 µs) in nine normal-hearing listeners. Explanations for the variability in the current study include small differences in the criterion used in the manual sensation level measurement, attention and fatigue, and more consistently good performance had extended training been used (Wright and Fitzgerald, 2001). Best and Swaminathan (2019, Fig. 2) also reported that ITD thresholds at 750 Hz did not correlate with the audiometric threshold at 750 Hz (or 350 and 1150 Hz). In the current study, ITD discrimination sensitivity similarly did not correlate with audiometric thresholds at 500 or 1000 Hz (*p*>0.05 for both).

Future behavioral and physiological experiments that examine ITD sensitivity as a function of frequency and level are required to better understand the properties of the neurons that convey fine-structure ITD information. Such experiments include animal studies in mammalian models to determine if they have a single dominant fine-structure ITD channel and its location in frequency. For example, primates may have a dominant channel near the same frequency region as humans but in animals with smaller heads (e.g., cats, ferrets, and chinchillas), it could be higher, if it exists at all (Verschooten et al., 2019). In addition, other binaural phenomena would likely also benefit from the joint exploration of frequency and level. For example, binaural masking level differences (improved detection of a tone in noise due to different interaural configurations of tone and noise) and the existence region of binaural pitches like Huggins pitch (when otherwise diotic white noise has a narrow band where the interaural phase progressively changes a full 360° cycle) also shows a precipitous drop near 1500 Hz (Culling, 1999). Mechanisms that require temporal fine structure ITD and IPD processing should show decreasing upper frequency limits as the level decreases. Divergence of effects for an upper frequency would suggest other mechanisms involved in the binaural phenomena, such as the use of envelope-based (ILD) cues (Goupell and Hartmann, 2006).

The proposed “dominant channel theory,” having only a single narrow band of ITD sensitive neurons near 700 Hz, is starkly different than the classic binaural display proposed by Jeffress (Jeffress, 1948; Stern et al., 1988; Stern et al., 2019), which consists of a matrix of frequency- and ITD-tuned neurons. Such a reimagining of human ITD processing overlaps with past discussions about mammalian vs avian binaural-processing models, the distribution and length of neural delay lines, and how the binaural system processes physiologically plausible and implausible ITDs (Thompson et al., 2006; Grothe et al., 2010; Stern et al., 2019; Yin et al., 2019; Litovsky et al., 2021). This dominant channel theory can be formulated to be consistent with the idea that the binaural system evolved in a manner to emphasize ecologically valid range of ITDs that provide unambiguous location information (Hartmann and Macaulay, 2014), helping to reconcile the discrepant interpretation of data from several past binaural-hearing studies.

In summary, the rapid 90 dB/octave rate of decline in fine-structure ITD sensitivity can, and arguably best, be explained by peripheral cochlear filtering. This suggests that listeners are performing off-frequency listening to neurons tuned near 700 Hz, possibly through a single dominant channel. Such a model contradicts the classic binaural display that utilizes a matrix of neurons that yield maximum firing rates at their best frequency and best interaural time difference.

## ACKNOWLEDGMENTS

We thank Taylor Beinke for assistance with data collection. We thank Jonas Klug and Mathias Dietz for helpful discussions concerning the frequency dependence of ITD discrimination.

## AUTHOR CONTRIBUTIONS

Conceptualization, methodology, software, investigation, data visualization, formal analysis, writing – original draft, funding acquisition, M.J.G.

Conceptualization, methodology, software, writing – review & editing, funding acquisition, G.C.S.

Methodology, investigation, writing – review & editing, B.T.W. Methodology, investigation, writing – review & editing, A.B.

Conceptualization, methodology, investigation, data analysis, writing – review & editing, funding acquisition, D.J.T.

## STATEMENTS AND DECLARATIONS

The authors declare no competing interests.

